# Implementation and Evaluation of Marker Rescue and qPCR Assays for Detection of Replication-Competent Retrovirus in Gene Therapy Products

**DOI:** 10.64898/2026.10.27.762168

**Authors:** Hamidreza Ebrahimiyan, Ali Sayadmanesh, Hossein Baharvand, Mohsen Basiri

## Abstract

Quality control of retroviral gene therapy vectors requires reliable detection of replication-competent retrovirus (RCR) particles. Cell-based assays remain the standard for detecting RCR, while PCR-based assays serve as rapid, complementary methods. Despite their widespread use, few studies provide comprehensive protocols and performance data to guide assay setup. Here we provide a detailed description and performance assessment of a cell-based firefly luciferase marker rescue assay for RCR testing. Additionally, we describe the setup and assessment of a qPCR-based RCR assay targeting retroviral *pol* gene sequence. Different viral copy numbers of MuLV were used to determine the final sensitivity of the marker rescue assay with and without initial amplification. Different q-PCR strategies were investigated for RCR testing using different types of primers, including env and pol genes. The marker rescue assay detected 73.2 RCR genome particles without amplification, while the amplification phase enabled detection of functional RCR particles and increased sensitivity, with the amplification step increasing viral particles by up to 5.23 × 10³ fold. The PCR-based RCR assay using retroviral *pol* gene could detect approximately 7.32 × 10^3^ viral particles. This study offers a resource for establishing RCR testing procedures for clinical-grade retroviral vector products.

## 1 Introduction

Various advancements have been made in the field of gene therapy that have expanded safety testing strategies in viral gene delivery methods. Retroviral and lentiviral vectors were developed by replacing helper genes, including group-specific antigen (gag), polymerase (pol), and envelope (env), with an exogenous gene of interest (vector) (1). These vectors are transferred to transient packaging cell lines (HEK-293T) to transiently express these genes alongside the transgene-carrying vector (2). In 2016, retroviral gene therapy was first granted authorization in Europe for adenosine deaminase deficiency (ADA) (3); the following year, retroviral gene therapy targeting non-Hodgkin’s lymphoma was similarly approved in the United States (4).

Following a span of three decades since the first approved gene therapy (5), no adverse events associated with retroviral gene transfer have been reported. The main concern in viral gene therapy is the development of Replication-Competent Retrovirus or lentivirus (RCR/L) particles (6, 7). While viral gene delivery systems are designed not to replicate, recombination events between the vector used for virus production and endogenous helper genes in the packaging cell lines, or contamination, could potentially generate an RCR/L during the manufacturing process or in patients after infusion (8–10). Reducing the homology sequence between the packaging sequence and vectors that was applied in the second generation of packaging cell lines, as well as segregating packaging genes into different plasmids as seen in the third generation such as PG13, has decreased the probability of RCR/L production in packaging cell lines (2). PG13 cells were derived from TK-NIH/3T3 cells which is based on the gibbon ape Leukaemia Virus (GaLV) envelope and murine leukemia viral (MuLV) gag- polymerase (11, 12).

RCR has demonstrated pathogenic potential in mice and non-human primates, causing malignancy (13, 14). Therefore, the Food and Drug Administration (FDA) requires vector products to be screened for RCR/L (15). The FDA recommends RCR testing by transfecting vector products into permissive cell lines and passaging them at least five times over three weeks (16). In the detection phase, cell-free media can be analyzed using methods such as marker rescue or colony-forming unit assays (17). RCR analysis is required on packaging cell lines and wherever retroviral particles are used in therapy that is approved for retroviral vector production in good manufacturing practices (GMP) (18). Cell-based RCR testing must be conducted for master or any working cell banks (MCB, WCB) and vector-producing cells (VPC) for 1% of banking or a maximum of 10^8^ cells (16). Five percent of the final vector product and MCB/WCB media must also be screened for any RCR. The FDA recommends detecting less than one RCR per dose in cell-based RCR testing. In addition to cell-based RCR testing, PCR-based methods are also recommended by the FDA for lot release testing of ex vivo transduced cells in time-limited situations (16). Using PCR techniques, patients undergoing gene therapy cell products, such as CAR T-cell therapy, should be monitored for RCR pre-treatment at 3, 6, and 12 months using PCR methods (16, 19, 20). Both cell media and cells in final products should be monitored in cell-based RCR testing. RCR testing screens for any RCR based on the tropism of the parental virus used to generate the vector (16).

Herein, retroviral packaging cell lines based on the HEK-293T cell line were used for retroviral transient production (21). The Platinum-A (Plat-A) cell line was developed to produce amphotropic MuLV envelope-based ecotropic MuLV retrovirus, optimized for amphotropic envelope (22). The MuLV-based retroviral vector was used for transduction of the PG13 producer cell line, which produces amphotropic GaLV envelope retrovirus (CAR PG13) (2). RCR testing with a marker rescue assay using Firefly luciferase was developed on the HEK- 293T cell line, a permissive cell line for both amphotropic MuLV and GaLV CAR retrovirus. The sensitivity of the marker rescue assay was assessed in different phases of testing to characterize the sensitivity of the test (23). RCR testing with PCR methods was also designed to amplify the pol gene instead of env to detect most producer cell lines, including those producing PG13 retroviral vectors, alongside Plat-E/A or most RCR control positive viruses such as VR-1450 with the Mo/A-MuLV envelope. The sensitivity of the PCR method was compared to the RCR rescue assay test.

## 2 Methods and Materials

### 2.1 Vector Generation

The Firefly luciferase coding sequence (sourced from the Royan Plasmid Bank) was cloned into the SFG.CNb30_opt.IRES.eGFP plasmid (Addgene plasmid # 22493) using traditional cloning methods (PCR amplification, restriction digestion, and ligation). The SFG.Luci.IRES.eGFP plasmid was utilized as a sensitive marker in the rescue assay test. Plasmid DNA was prepared using the QIAGEN Plasmid Maxi Extraction Kit (USA, cat: 12163).

### 2.2 Cell Lines and Culture Conditions

Plat-A, Plat-E, HEK-293T cell lines sourced from the Royan Cell Bank, and PG13 (ATCC CRL-10686) were used as cell sources. The single positive CAR PG13 cell line producer was sourced from the Royan Cell Bank. All cells were cultured in IMDM (Iscove’s Modified Dulbecco’s Medium, Gibco cat: 21980032) with 10% Fetal Bovine Serum (Characterized FBS, HyClone cat: SH30071.03IH30-45) and 2 mM GlutaMax (100x, Gibco cat: 35050-061) and incubated at 37°C in 5% CO2.

### 2.3 Transfection and Transduction

For Plat-E/A transfection, a 100 mm treated cell culture plate was treated with 0.1% gelatin for 1 hour at 37°C. After overnight seeding of 4.5×10^6^ cells, they were transfected with the vector of interest using Lipofectamine 3000 (Invitrogen, Part no. 100022234) according to the manufacturer’s instructions. The media was renewed the next day, and virus collection and snap freezing were performed 24 and 48 hours after transfection. The media containing the viruses were stored at -80°C for HEK-293T cell transduction. In a 24-well plate, 4×10^4^ cells were seeded for HEK-293T transduction. Virus supernatants containing 10 μg/ml polybrene (MERK, SKU: TR-1003-G) were added to each well. The media was changed the next day, and the re-transduction protocol was applied when confluency reached 80%. The transduction procedure was repeated until 80% of HEK-293T cells were GFP+ (GFP positive). The sharp positive single clone of HEK-293T was sorted based on GFP to be utilized for the rescue assay test in the detection phase.

### 2.4 Amplification Phase in Rescue Assay Test

On day 0, HEK-293T cells (up to the 10th passage) were seeded in 24-well plates at a concentration of 1×10^5^ cells per well. On day 1, during the transduction stage, the media was discarded, and 1 ml of sample with 10 μg/ml polybrene was added. On the second day, after overnight culture, the media was refreshed with 1 ml of new media. On day 3, all wells were split at the same concentration as on day 0. Every 2 days, the media was changed, and based on cell confluences (80-90%), splitting was done every 3 or 4 days after the last split. After 5 passages, the cells were seeded with 2×10^5^ cells in each well. The next day, the media was changed, and after 24 hours, it was harvested and filtered (0.45 µm) for use in the detection phase.

### 2.5 Detection Phase in Rescue Assay Test

Stage 1: On day 0, HEK-293T Luci^+^ cells were seeded in 24-well plates at a concentration of 1×10^5^ cells per well. On day 1, the media was discarded, and 1 ml of amplified sample with 10 ug/ml polybrene was added. On day 2, after overnight, the media was renewed. On day 3, all wells were split at the same concentration as on day 0. Every 2 days, the media was changed, and based on cell confluences (80-90%), splitting was done every 3 or 4 days after the last split. Following two passages, 2×10^5^ cells were seeded into each well. The media was renewed the following day, and the supernatant was harvested and filtered (0.45 µm) for use in stage 2 of the detection phase after 24 hours. Stage 2: All steps are similar to stage 1, except using HEK- 293T instead of HEK-293T Luci^+^. After the second passage, when cell confluences were at 80- 90%, the cells were trypsinized and washed for luciferase reaction detection with Firefly Luciferase HTS Assay (cat: # SCT150) according to the manufacturer’s protocol.

### 2.6 RCR Positive Control Production

The MuLV virus obtained from ATCC (VR-1450) was used as the RCR control positive. One sample of VR-1450 was amplified in the amplification phase and aliquoted into several vials. Each vial was titrated using the Retro-X™ Q-PCR Titration Kit (cat: 631453) according to the manufacturer’s protocol. The detected copy numbers were calculated based on the standard curve linear formula.

Detected copy number (copies/μl) = 10^(Ct−b)/a^ The original concentration of virus (copies/ml)

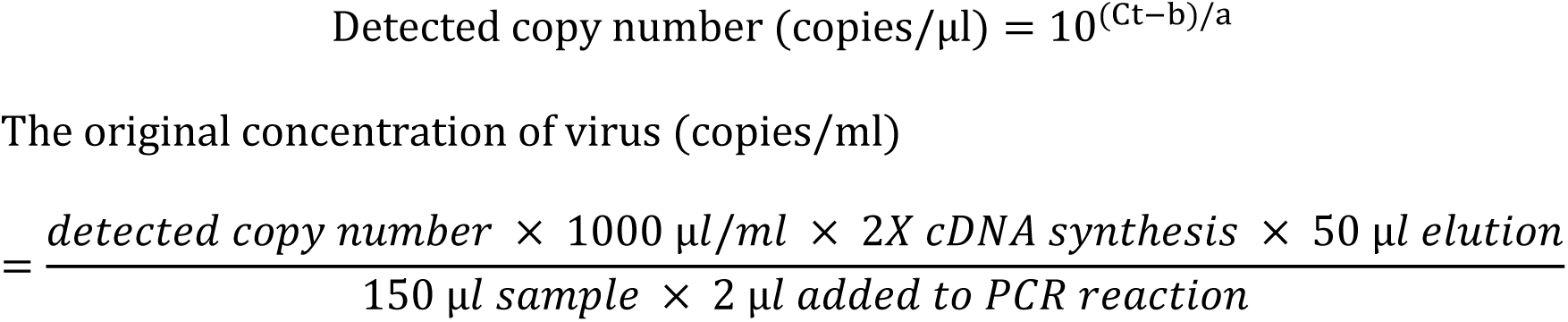

### 2.7 Sensitivity of rescue assay test without the detection phase

The MuLV aliquots were diluted with serial dilutions (10^-2^ to 10^-5^). Only the amplification phase was applied to determine the sensitivity of the rescue assay test without the detection phase. After the amplification phase, viral titrations were applied using the Retro-X™ Q-PCR titration kit. The detected copy numbers were calculated based on the standard curve linear formula.

### 2.8 Sensitivity of rescue assay test without amplification phase

The MuLV aliquots were diluted at two different times with serial dilutions (1⁄2, 1⁄4, 1⁄8, 10^-1^ to 10^-7^). Only the detection phase was applied to determine the sensitivity of the rescue assay test without the amplification phase.

### 2.9 Sensitivity of rescue assay test with amplification phase

To determine the sensitivity, the MuLV was diluted from 10^-1^ to 10^-7^, and both phases were conducted. The viral copy numbers were applied in two repeats, and serial dilutions were performed to 0.62 copy numbers. Additionally, 3, 2, and 1 viral copy numbers were applied in different repeats for the rescue assay test with the amplification phase.

### 2.10 PCR molecular assay using envelop and polymerase primers

Two different envelopes were targeted for primer design to detect the MuLV or GaLV envelope gene in any samples. The polymerase primer was also used to find any polymerase gene in different samples. The sequence of primers is listed in Table 4. Three different cell lines, including Plat-A/E and PG13, were used as samples. DNA was extracted using the QIAamp DNA Micro Kit (cat: 56304) according to the manufacturer’s protocol. The reaction mixture comprised 10 μl of PCR Master Mix, 2 μl of cDNA template, 1 μl of each Pol primer, and ribonuclease-free water, culminating in a total volume of 20 μl. The PCR conditions were established as follows: 95°C for 15 min, 40 cycles of 95°C for 30 seconds, 60°C (Pol) or 62°C (Envelopes) for 1 min, 72°C for 1 min, and then 72°C for 15 min.

### 2.11 Sensitivity of RCR-PCR with polymerase primer

The MuLV aliquots at two different times were diluted with serial dilutions (10^-2^ to 10^-5^). After that each sample performed 2 hours centrifugation at 14000 g. Pellet from the resultant was re- suspended in 140 ul, of phosphate-buffered saline (PBS). Viral RNA was extracted using the QIAamp® Viral RNA Mini (cat: 52904) according to the manufacturer’s protocol. Viral cDNA was synthesized immediately using the QuantiTect® Reverse Transcription (cat: 205311) following the company’s protocol. Conventional PCR was conducted using Taq DNA Polymerase 2x Master Mix RED (Cat no. A190307, Ampliqon) and a conventional PCR system under the same conditions as for DNA samples.

### 2.12 Statistical analysis

For data calculation, SPSS software v.29 (IBM, Armonk, NY) was used. The linear regression test was applied, and *Pv* < 0.05 and R_2_ > 0.98 were defined as a statistically significant level. For plotting, GraphPad Prism version 10 software (GraphPad Software, La Jolla, CA, www.graphpad.com) was used. Mean ± STD was expressed as scale data, and *Pv* < 0.05 was defined as a statistically significant level. ImageJ software V18 was used to quantify luciferase signals. The quantified results were normalized based on the control positive, which was applied as 100.

## 3. Results

### 3.1 MuLV virus titration

For the MuLV virus aliquots, the VR-1450 was amplified in the amplification phase of the rescue assay test (Figure 1). The MuLV aliquot titration was conducted. The amplification plot for standards was shown in Figure 2A. The standard curve demonstrates a strong linear regression between the Ct values and the RNA copy numbers (log scale) with R_2_ = 0.9994, PCR efficiency of 99.914%, and regression *Pv*< 0.001 (Figure 2C). The calculated original concentration was 1.30×10^8^ for undiluted MuLV and 1.63×10^7^ for a 10-fold dilution of MuLV. The average of both copy numbers was applied as the real concentration (7.32×10^7^) (Table 1).

**FIGURE 1:**
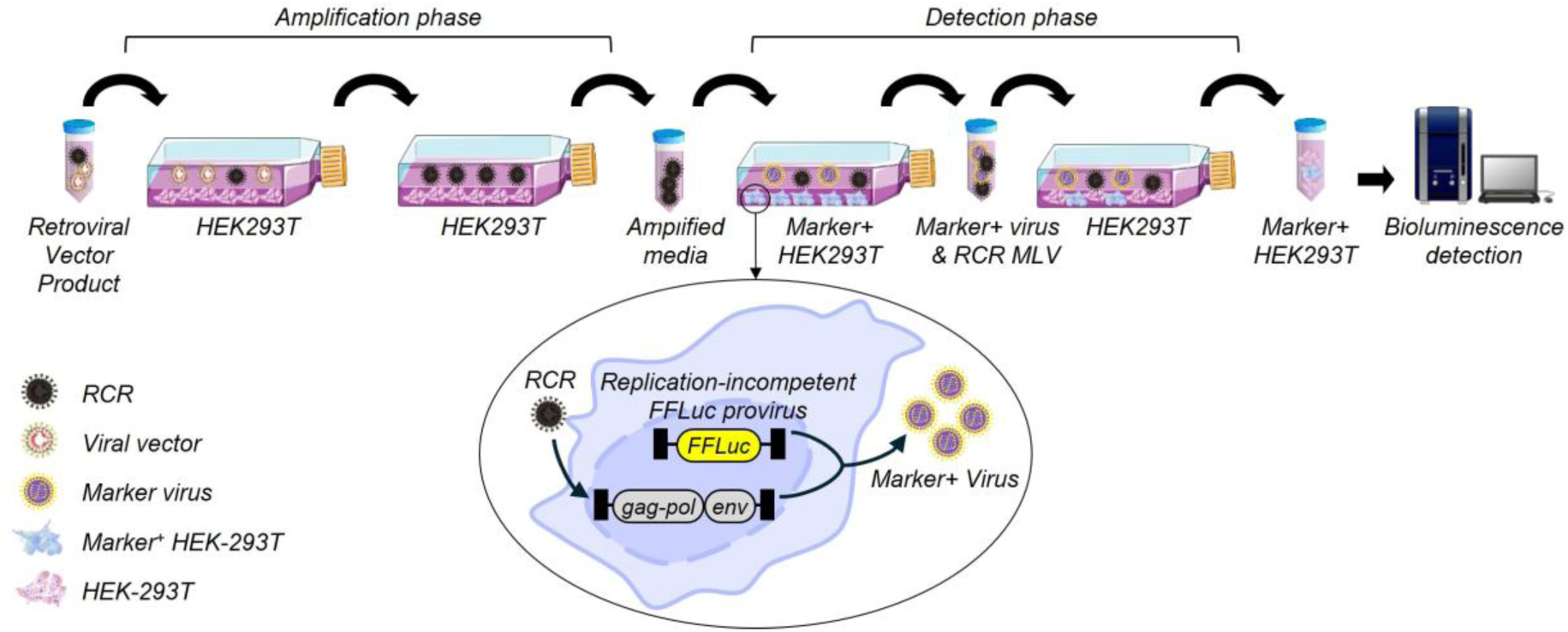
RCR rescue assay test. Amplification phase: potential RCR particles in viral vector supernatants are allowed to amplify in a permissive cell line within 3-weeks to increase the titer of RCR to be easily detectible. Detection phase: If RCR is present, it will rescue the marker and could be detected with related tools.

**FIGURE 2:**
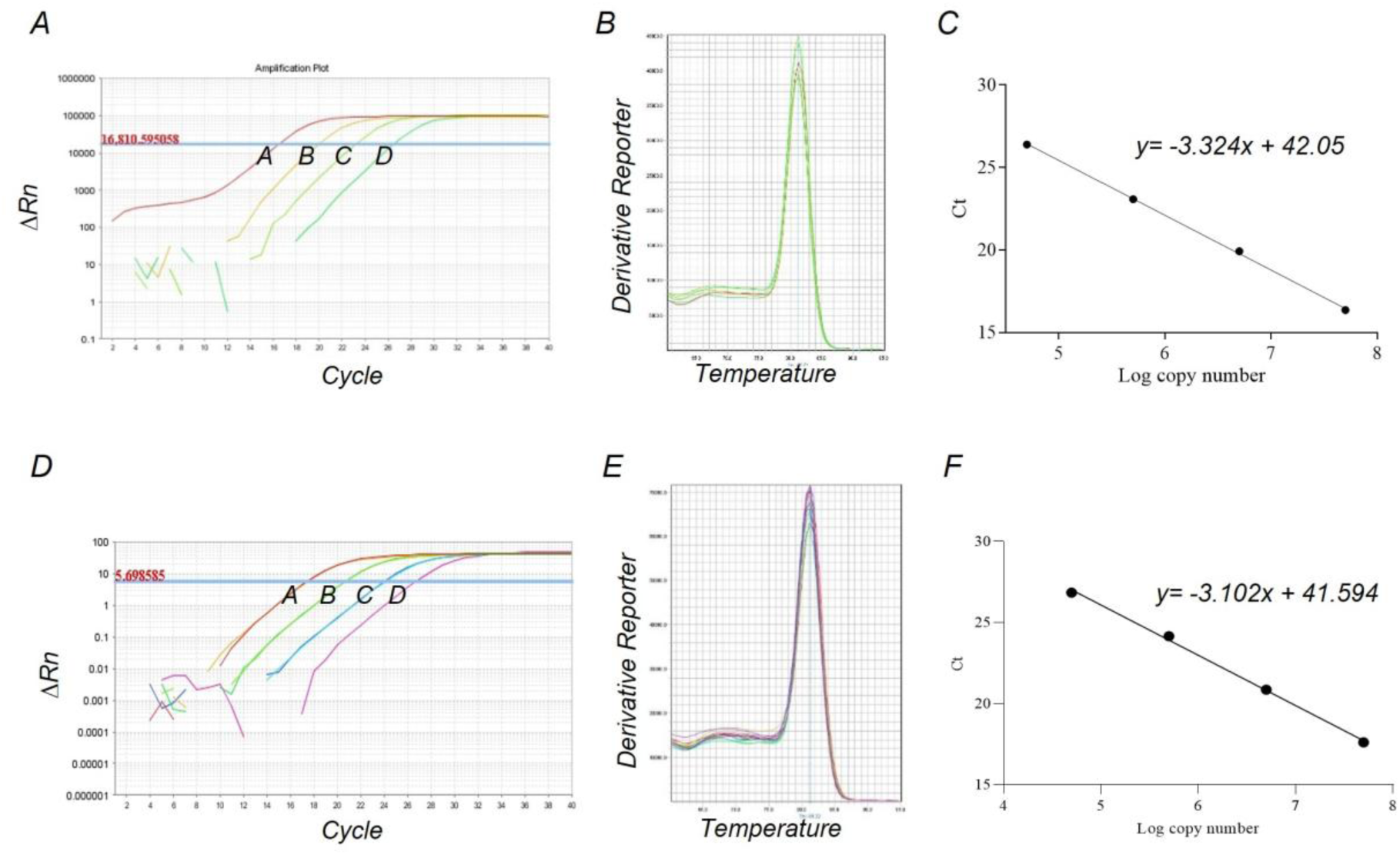
The MuLV titration before and after the amplification phase in the rescue assay test. **A.** Amplification plot, Standard copy number dilution: A: 5×10^7^, B: 5×10^6^, C: 5×10^5^, D: 5×10^4^. **B.** Melt curve plot of standards and samples. **C.** A standard curve created from panel A amplification plot demonstrates a strong linear regression between the Ct values and the RNA copy numbers (log scale). **D.** Amplification plot, Standard copy number dilution: A: 5×10^7^, B: 5×10^6^, C: 5×10^5^, D: 5×10^4^. **E.** Melt curve plot of standards and samples. **F.** A standard curve created from panel D amplification plot demonstrates a strong linear regression between the Ct values and the RNA copy numbers (log scale).

**Table 1:** MuLV titration

| Initial sample | Dilution factor | Absolute copy number (copies/μl) | Q-PCR CT average | Detected copy number (copies/μl) | Original concentration* (copies/ml) |
| --- | --- | --- | --- | --- | --- |
| Standard | $10^{-1}$ | $5 \times 10^7$ | 16.37 | $5.32 \times 10^7$ | NA |
| Standard | $10^{-2}$ | $5 \times 10^6$ | 19.93 | $4.51 \times 10^6$ | NA |
| Standard | $10^{-3}$ | $5 \times 10^5$ | 23.08 | $5.09 \times 10^5$ | NA |
| Standard | $10^{-4}$ | $5 \times 10^4$ | 26.4 | $5.11 \times 10^4$ | NA |
| MuLv | 1 | NA | 23.47 | $3.89 \times 10^5$ | $1.30 \times 10^8$ |
| MuLv | $10^{-1}$ | NA | 26.46 | $4.90 \times 10^4$ | $1.63 \times 10^7$ |

MuLv, Murine leukemia virus; NA, Not Applicable. \*: The concentration of virus in main sample.

### 3.2 Amplification of MuLV up to the saturation concentration

The MuLV-diluted samples ranging from 10^-2^ to 10^-5^ were applied to define the amplification ratio of the amplification phase in the rescue assay test. The amplification plot for standards was shown in Figure 2D. A standard curve demonstrates a strong linear regression between the Ct values and the RNA copy numbers (log scale), with R_2_ = 0.9978, PCR efficiency of 110.0%, and regression *Pv*: 0.001 (Figure 2F). The calculated original concentrations were 3.59×10^7^, 7.86×10^6^, 8.53×10^6^, and 3.83×10^6^, respectively. The amplification ratio ranges from 5.23×10^3^ to 49.04, depending on the starting titer of the virus (Table 2). In the amplification phase of the rescue assay test, any functional MuLV could be amplified until saturation concentration, which could be easily detected in the detection phase.

**Table 2:**
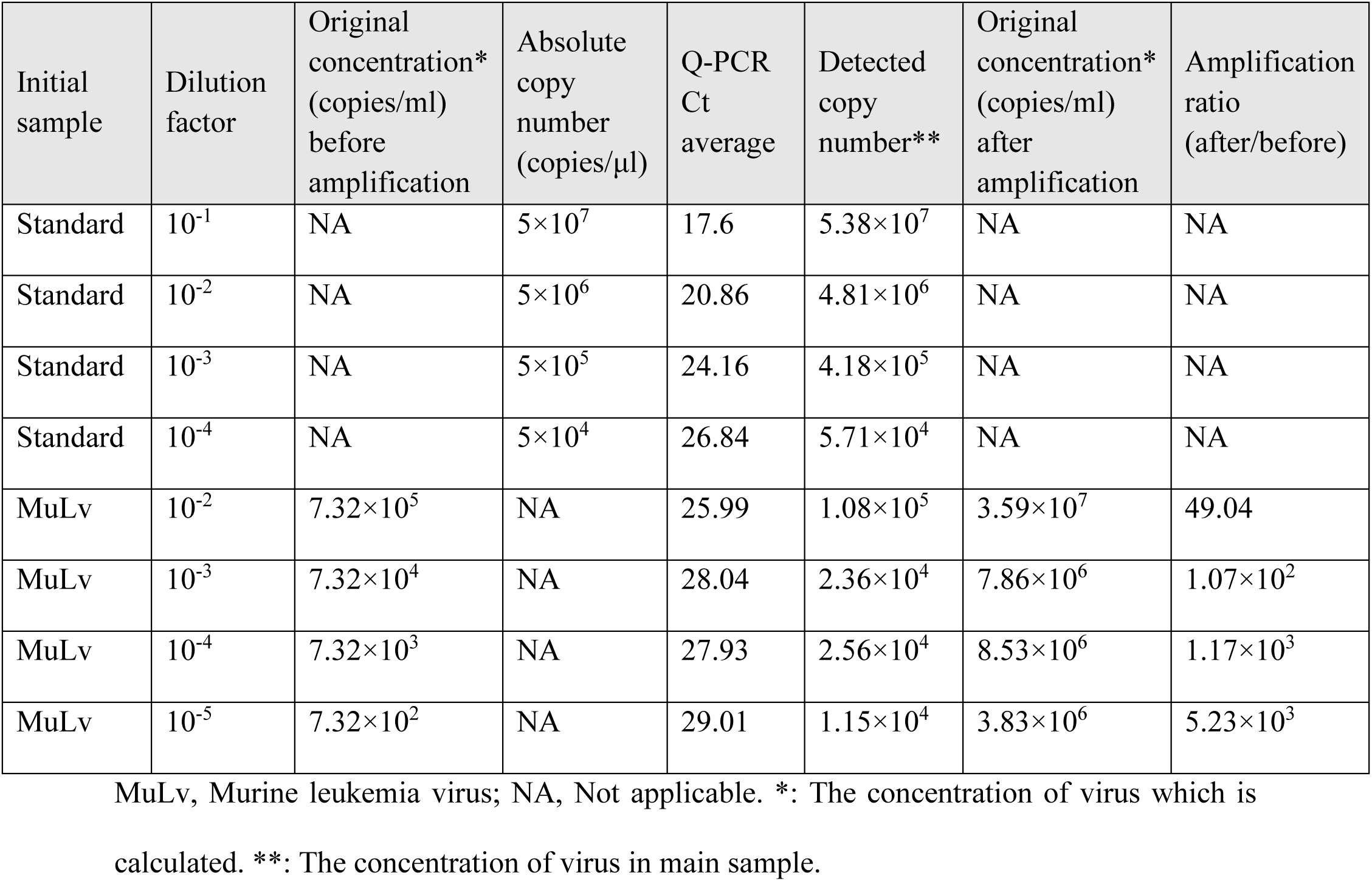
MuLV titration after amplification phase

### 3.3 Sensitivity of reporter rescue assay without initial amplification step

The MuLV dilution ranges from 1, 1⁄2, 1⁄4, 1⁄4, 10^-1^ to 10^-7^ were applied to define the sensitivity of the detection phase in the rescue assay test. The MuLV dilution range from 1 to 10^-6^ was positive in this condition. The rescue assay test without the amplification phase (which resulted in more viral particles) could also detect 73.2 copies of viruses (Figure 3A, Table 3). The quantified normalized results revealed a strong regression between the original concentrations of diluted MuLV samples (*Pv*: 0.006) (Figure 3B). Herein, we found that the detection phase is able to detect only 73.2 viral copy numbers in the sample.

**FIGURE 3:**
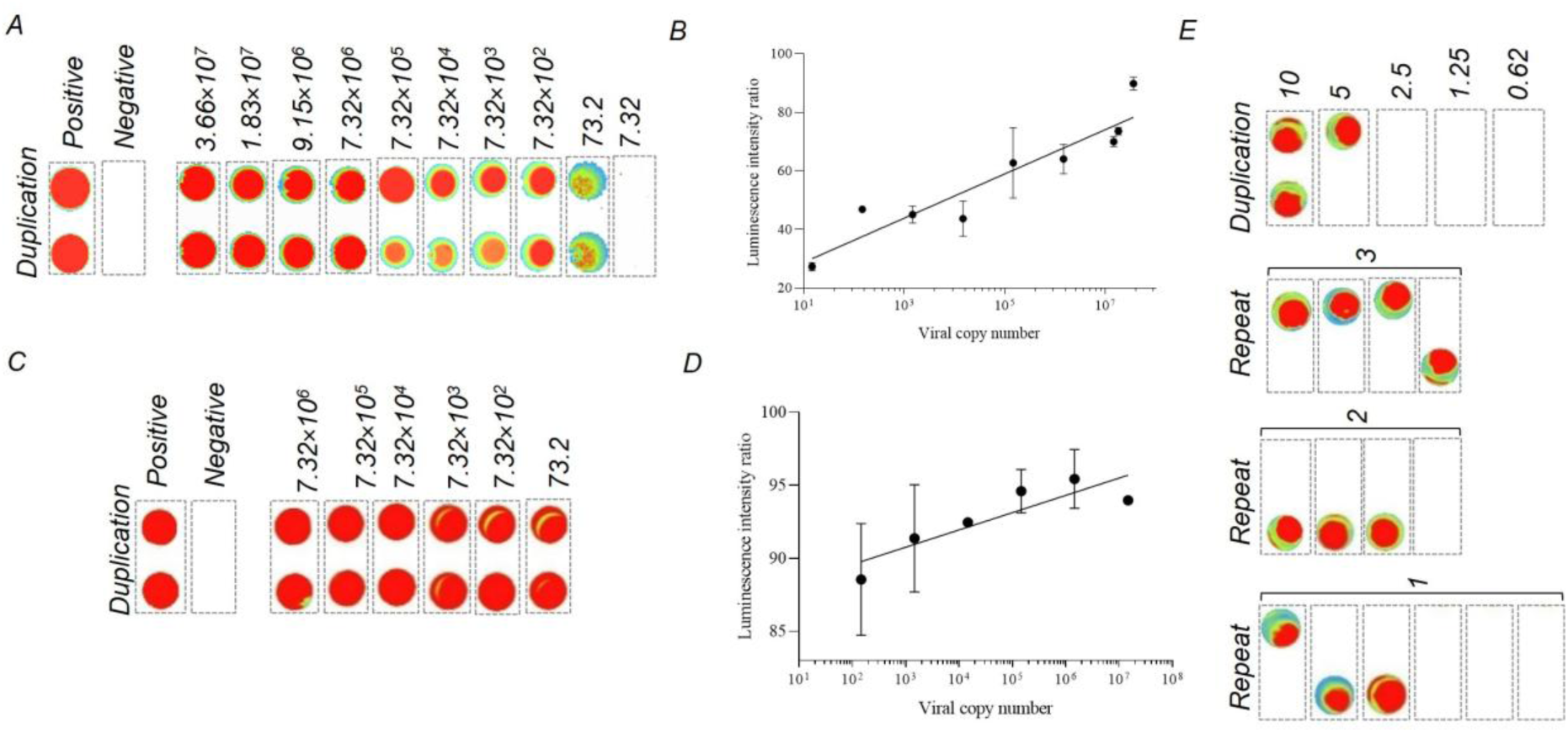
The rescue assay test with or without amplification phase. **A.** Rescue assay results without amplification phase with different titers of MuLV (the titer is the number of viruses before the test). **B.** A standard curve created from panel A results, which is quantified with ImageJ software. The positive control (with 7.32×10^7^ titer) quantitative results were applied as 100. The negative control (media only) quantitative results were applied as 0. The mean and SD of duplicated samples were applied. **C.** Rescue assay results with amplification phase with different titers of MuLV (the titer is the number of viruses before the test). **D.** A standard curve created from panel C results, which is quantified with ImageJ software. The positive control (with 7.32×10^7^ titer) quantitative results were applied as 100. The negative control (media only) quantitative results were applied as 0. The mean and SD of duplicated samples were applied. **E.** Rescue assay results with amplification phase with low copy numbers of MuLV.

**Table 3:** Results summary

| MuLV dilution | MuLV original concentration (copy/ml) | Molecular RCR (PCR) | Rescue assay results without amplification | Rescue assay results with amplification | MuLV original concentration after amplification phase* |
| --- | --- | --- | --- | --- | --- |
| 1 | $7.32 \times 10^7$ | NA | Positive | Positive | NA |
| $1/2$ | $3.66 \times 10^7$ | NA | Positive | NA | NA |
| $1/4$ | $1.83 \times 10^7$ | NA | Positive | NA | NA |
| $1/8$ | $9.15 \times 10^6$ | NA | Positive | NA | NA |
| $10^{-1}$ | $7.32 \times 10^6$ | NA | Positive | Positive | NA |
| $10^{-2}$ | $7.32 \times 10^5$ | Positive | Positive | Positive | $3.59 \times 10^7$ |
| $10^{-3}$ | $7.32 \times 10^4$ | Positive | Positive | Positive | $7.86 \times 10^6$ |
| $10^{-4}$ | $7.32 \times 10^3$ | Positive | Positive | Positive | $8.53 \times 10^6$ |
| $10^{-5}$ | $7.32 \times 10^2$ | Positive/<br>Negative | Positive | Positive | $3.83 \times 10^6$ |
| $10^{-6}$ | 73.2 | Negative | Positive | Positive | NA |
| $10^{-7}$ | 7.32 | Negative | Negative | Positive | NA |

NA, Not applicable, \*: The concentration of virus in main sample.

**Table 4:**
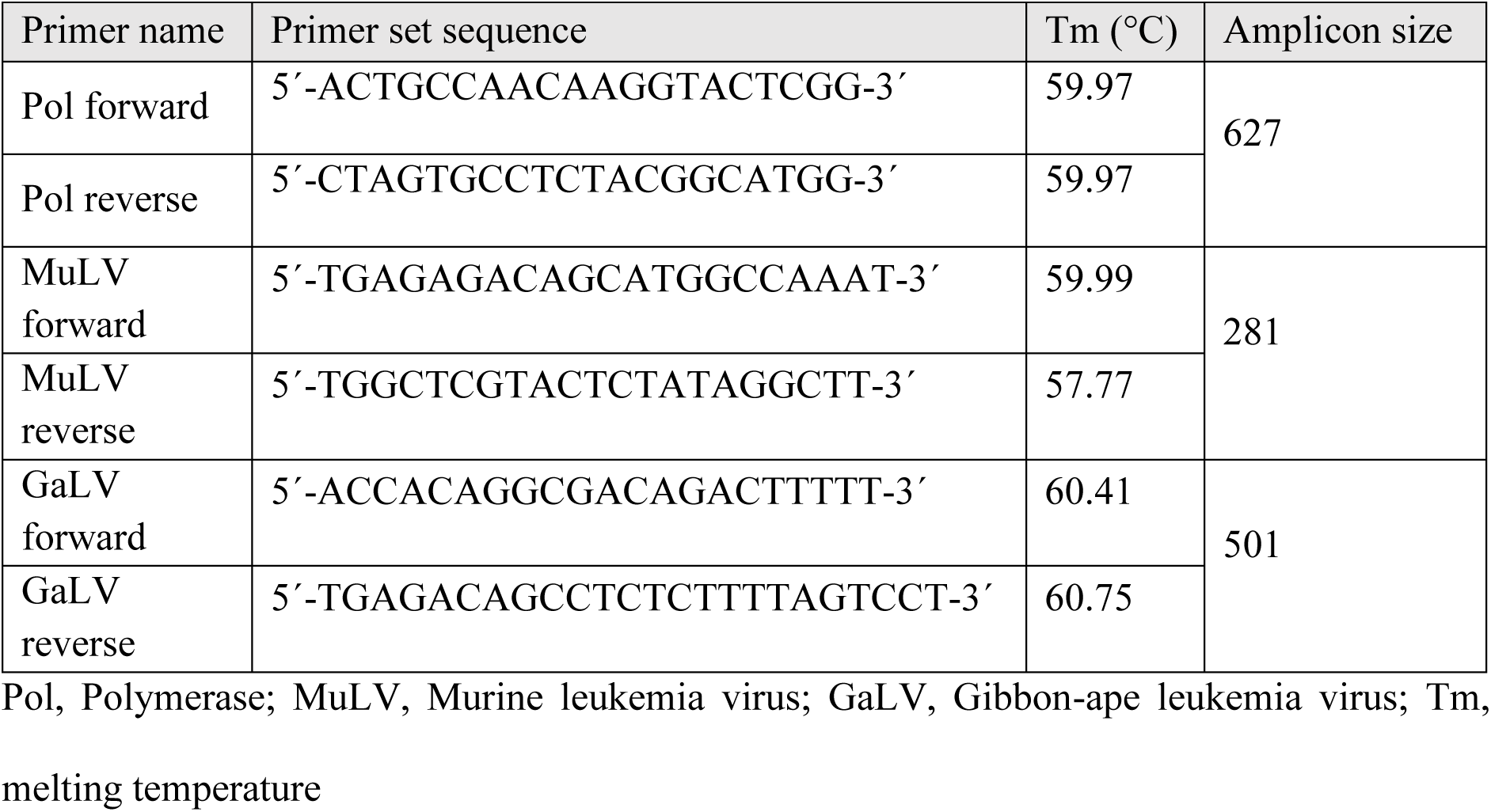
Details of primers

### 3.4 High sensitivity of reporter rescue assay with initial amplification step

The MuLV dilution ranges from 10^-1^ to 10^-6^ were applied to define the sensitivity of the rescue assay test results with amplification. The positive results were detected in all samples (Figure 3C). The quantified normalized results were shown in Figure 3D. Viral particles in 10, 5 were also positive in two repeats. 3 viral copy numbers were positive 4 times with 8 repeats (50%). 2 viral copy numbers were positive 3 times with 8 repeats (37.5%). 1 viral particle was positive times in 12 repeats (25%) (Figure 3E, Table 3). However, the starting titer of viruses was based on average molecular Q-PCR results, which means any genetic particles were tittered; interestingly, only one MuLV viral particle was also marked as positive in the rescue assay test.

### 3.5 Validation of PCR-based RCR assay across different packaging cell lines

The MuLV primer could detect the MuLV envelope in the Plat-A/E cell line; however, there was not a completely identical sequence. The PG13 cell line could not be detected with this primer (Figure 4A). The GaLV primer only detected the GaLV envelope in PG13, and Plat- A/E could not be detected with this primer (Figure 4B). The Pol primer, which detected the polymerase gene, could detect all three cell lines and any RCR that may develop from these producer cell lines (Figure 4A). Herein, we developed a new set of primers for RCR-PCR testing that were not dependent on envelopes and could detect RCR in most producer cell lines.

**FIGURE 4:**
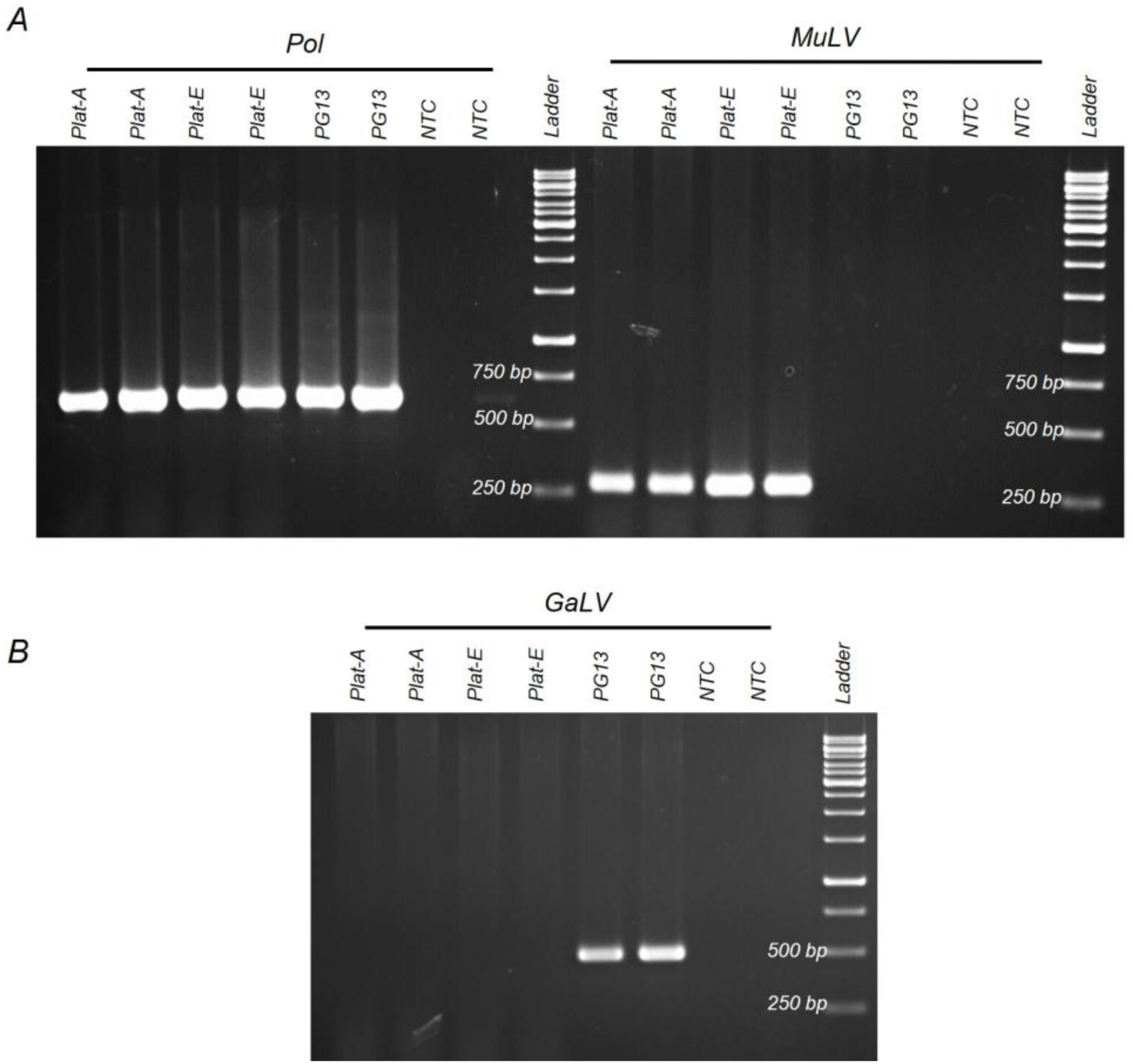
Different primer in molecular RCR. **A.** Polymerase and MuLV-related primers in three different cell lines. **B.** GaLV-related primer in three different cell lines. NTC: no template control.

**FIGURE 5:**
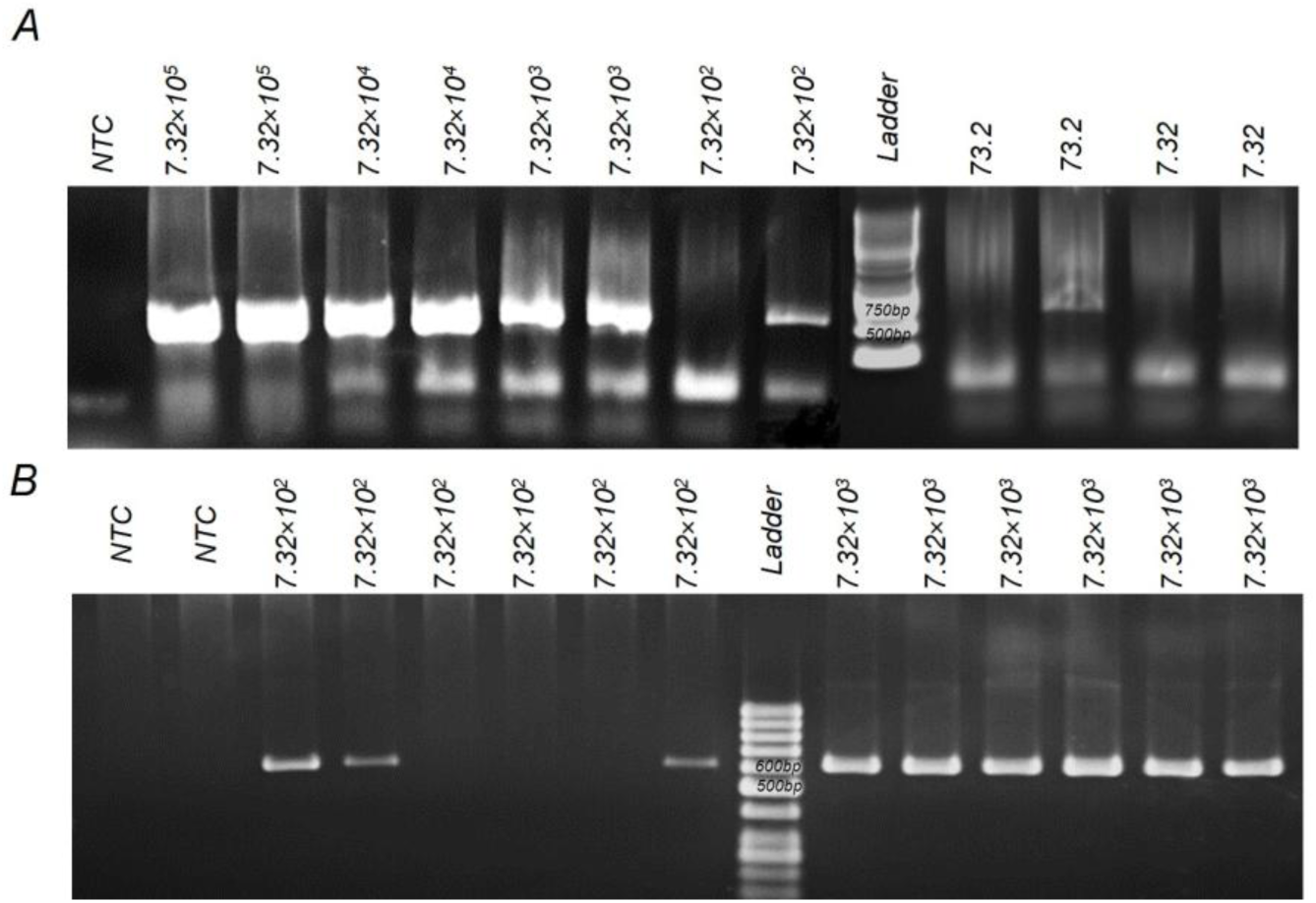
The sensitivity of pol PCR to detect different MLV virus titer. **A.** Different MLV titers were applied. **B.** The only two titers of MLV (7.32×10^3^ and 7.32×10^2^) virus were applied to find detection sensitivity. The 7.32×10^3^ viral titers were the detection limit in the PCR test. NTC: no template control.

### 3.6 Sensitivity of PCR-based RCR assay

The MuLV dilution ranges from 10^-2^ to 10^-7^ were applied to define the sensitivity of the detection phase in RCR-PCR testing. Based on the results, the PCR method could detect 7.32×10^3^ RCR particles in all repeated samples; however, the 7.32×10^2^ RCR particles could be detected in half of the repeats (50%). The sensitivity of the molecular test is less than the cell-based strategy, but it takes only one day to report the first results.

## 4. Discussion

Although new generations of gene therapy vectors and producer cell lines reduce the risk of RCR formation, screening of gene therapy products remains an essential step in product release quality controls. Malignancy is a major concern in RCR infection, as shown in mice and non-human primates (13, 14). The US FDA recommends RCR testing in three steps: first, RCR testing in release criteria for any retroviral vector lots; second, monitoring patients who received retroviral-related products; and third, RCR testing in any retro-transduced cell products cultured ex vivo for more than 4 days (16, 17). Screening for RCR using cellular methods remains the gold standard test, capable of identifying <1 RCR dose in samples (16). For cell-based RCR, commonly used tests include the marker rescue assay and colony-forming units (17, 24–26). The colony-forming unit method requires a professional staff to screen the entire plate to detect any colonies under a microscope, which could be problematic for releasing GMP products. However, in the marker rescue assay, any signals detected can be analyzed with advanced tools by non-professionals. In the marker rescue assay, enzymatic reactions also amplify the sensitivity of signals (16, 27). Cellular RCR testing is time-consuming, limiting its applicability in situations like product release for gene therapy. Molecular methods could be useful for RCR testing in a few hours (27). The similarity between viral vectors and RCR poses difficulties in molecular detection, as both include viral genes encoding proteins such as integrase, reverse transcriptase, or capsid, making protein detection methods or reverse transcriptase assays unhelpful. Viral vectors that are replication-defective lack genetic material, including Gag, Pol, and Env genes present in vector producer cell lines. Detecting these genes with PCR is a main issue in developing RCR-PCR testing. Another weakness of this method is false positive results in direct producer cell media due to cellular free DNA or extracellular vesicles. Based on FDA guidelines (16), PCR methods for final products and patient follow-up could be used instead of cellular RCR testing. In this case, any potential RCR particles may have enough time to amplify and reach the detection limits of PCR; however, cellular RCR testing is also required to confirm the PCR results.

In the current study, Firefly luciferase was developed for cellular RCR testing, and the sensitivity of each phase and the entire rescue assay testing were analyzed. The rescue assay test without an amplification phase (used to amplify viruses) could have detected approximately 73.2 RCR copy number particles. However, the rescue assay test with an amplification phase could have detected any functional RCR particle in samples. It is important to note that these 73.2 particles were calculated based on q-PCR, which cannot distinguish between infectious and non-infectious viruses. The amplification phase of the test, used to increase sensitivity, revealed an amplification of 5.23×10^3^ fold of 7.32×10^2^ RCR particles. However, the 7.32×10^5^ RCR particles amplified in 49.04 folds due to viral saturation in the media. In the case of 73.2 viral particles, RCR marker rescue assay testing revealed a strong positive signal, similar to the highest titer in the marker rescue assay test without an amplification phase, establishing a high amplification ratio in the amplification phase. The amplification phase was sufficient to amplify any RCR particle to the detection limits of the detection phase in the marker rescue assay testing. In RCR-PCR, the challenge is developing a method that can be useful for different vector production methods, such as Plat or PG13. In some cases, producing viral vectors in different cell types and selecting the same helper gene instead of an envelope could be helpful in developing a general molecular test for RCR testing. We developed PCR-RCR testing based on a polymerase present in most cases of RCRs. In our experiment, PCR-RCR testing could detect approximately 7.32×10^3^ RCR particles, with 7.32×10^2^ particles also detected in half of the repeats.

## 5. Conclusion

In this study, we developed a marker rescue assay based on Firefly luciferase and finally checked the sensitivity of each phase of the marker rescue assay testing. The marker rescue assay test we performed was able to detect any RCR particles in the given sample, however the detection phase also identifying RCR viral particles at an approximate count of 73.2 particles. The amplification phase also amplified the viral particles by at least 5.23×10^3^ fold, increasing the sensitivity of the marker rescue assay testing. RCR-PCR methods could be applied for the release of final products, detecting approximately 7.32×10^3^ viral particles. This general PCR method could be useful for different producer banks in gene therapy treatment and is not limited to specific viral envelopes or strains.

## Author Contributions

Hamidreza Ebrahimiyan: Investigation, methodology, formal analysis, data curation and writing original draft. Ali Sayadmanesh: writing – review and editing and data curation. Mohsen Basiri: Conceptualization, writing – review and editing, data curation and supervision. Hossein Baharvand: funding acquisition, review, editing and supervision.

## Acknowledgement

All donors who facilitated the accomplishment of this study are deeply acknowledged. We thank all staff and technicians at the Royan Institute for Stem Cell Biology and Technology for their technical assistance.

## Data Availability Statement

The data generated and analyzed during the study are available as requested.

## Declaration of generative AI and AI-assisted technologies in the writing process

During the preparation of this work the authors used ChatGPT (OpenAI) in order to correct spelling and grammar errors and improve readability of the text. After using this tool, the authors reviewed and edited the content as needed and take full responsibility for the content of the published article.

## Reference

1. Ayala Ceja M, Khericha M, Harris CM, Puig-Saus C, Chen YY. CAR-T cell manufacturing: Major process parameters and next-generation strategies. Journal of Experimental Medicine. 2024;221(2):e20230903.

2. Coroadinha AS, Gama-Norton L, Amaral AI, Hauser H, Alves PM, Cruz PE. Production of retroviral vectors. Current gene therapy. 2010;10(6):456–73.

3. Aiuti A, Roncarolo MG, Naldini L. Gene therapy for ADA-SCID, the first marketing approval of an ex vivo gene therapy in Europe: paving the road for the next generation of advanced therapy medicinal products. EMBO Mol Med. 2017;9(6):737–40.

4. Ong MZ, Kimberly SA, Lee WH, Ling M, Lee M, Tan KW, et al. FDA-approved CAR T-cell Therapy: A Decade of Progress and Challenges. Curr Pharm Biotechnol. 2024;25(11):1377–93.

5. Blaese RM, Culver KW, Chang L, Anderson WF, Mullen C, Nienhuis A, et al. Treatment of severe combined immunodeficiency disease (SCID) due to adenosine deaminase deficiency with CD34+ selected autologous peripheral blood cells transduced with a human ADA gene. Amendment to clinical research project, Project 90-C-195, January 10, 1992. Hum Gene Ther. 1993;4(4):521–7.

6. Salazar-Fontana LI, Havert M. Current Regulatory Landscape for Gene Therapy Product Development and the Role of Biomarkers. Drug Development for Gene Therapy: Translational Biomarkers, Bioanalysis, and Companion Diagnostics. 2024:429–48.

7. Food U, Administration D. Testing for Replication Competent Retrovirus (RCR)/Lentivirus (RCL) in Retroviral and Lentiviral Vector Based Gene Therapy Products— Revisiting Current FDA Recommendations. 2010.

8. Cesana D, Ranzani M, Volpin M, Bartholomae C, Duros C, Artus A, et al. Uncovering and Dissecting the Genotoxicity of Self-inactivating Lentiviral Vectors In Vivo. Molecular Therapy. 2014;22(4):774–85.

9. Dull T, Zufferey R, Kelly M, Mandel RJ, Nguyen M, Trono D, et al. A third-generation lentivirus vector with a conditional packaging system. J Virol. 1998;72(11):8463–71.

10. Hacein-Bey-Abina S, Pai S-Y, Gaspar HB, Armant M, Berry CC, Blanche S, et al. A Modified γ-Retrovirus Vector for X-Linked Severe Combined Immunodeficiency. New England Journal of Medicine. 2014;371(15):1407–17.

11. Kochenderfer JN, Feldman SA, Zhao Y, Xu H, Black MA, Morgan RA, et al. Construction and preclinical evaluation of an anti-CD19 chimeric antigen receptor. Journal of immunotherapy. 2009;32(7):689–702.

12. Graber DJ, Cook WJ, Sentman M-L, Murad-Mabaera JM, Sentman CL. Human CD4+ CD25+ T cells expressing a chimeric antigen receptor against aberrant superoxide dismutase 1 trigger antigen-specific immunomodulation. Cytotherapy. 2024;26(2):126–35.

13. Cornetta K, Nguyen N, Morgan RA, Muenchau DD, Hartley JW, Blaese RM, et al. Infection of human cells with murine amphotropic replication-competent retroviruses. Human gene therapy. 1993;4(5):579–88.

14. Donahue R, Kessler S, Bodine D, McDonagh K, Dunbar C, Goodman S, et al. Helper virus induced T cell lymphoma in nonhuman primates after retroviral mediated gene transfer. The Journal of experimental medicine. 1992;176(4):1125–35.

15. Health UDo, Services H. Supplemental guidance on testing for replication-competent retrovirus in retroviral vector-based gene therapy products and during follow-up of patients in clinical trials using retroviral vectors. Human gene therapy. 2001;12(3):315–20.

16. Food U, Administration D. Testing of Retroviral Vector-Based Human Gene Therapy Products for Replication Competent Retrovirus During Product Manufacture and Patient Follow-up. 2020: 1–14. 2021.

17. Cornetta K, Wilson CA. 23 Safety of Retroviral Vectors: Regulatory and Technical Considerations. Concepts in genetic medicine. 2008:277.

18. Gee AP. Gmp car-T cell production. Best Practice & Research Clinical Haematology. 2018;31(2):126–34.

19. Sastry L, Cornetta K. Detection of replication competent retrovirus and lentivirus. Methods in molecular biology (Clifton, NJ). 2009;506:243–63.

20. Cornetta K, Yao J, House K, Duffy L, Adusumilli PS, Beyer R, et al. Replication competent retrovirus testing (RCR) in the National Gene Vector Biorepository: No evidence of RCR in 1,595 post-treatment peripheral blood samples obtained from 60 clinical trials. Molecular Therapy. 2023;31(3):801–9.

21. Sheet PD. Platinum-A Retroviral Packaging Cell Line, Amphotropic.

22. Morita S, Kojima T, Kitamura T. Plat-E: an efficient and stable system for transient packaging of retroviruses. Gene Therapy. 2000;7(12):1063–6.

23. Skrdlant LM, Armstrong RJ, Keidaisch BM, Lorente MF, DiGiusto DL. Detection of Replication Competent Lentivirus Using a qPCR Assay for VSV-G. Mol Ther Methods Clin Dev. 2018;8:1–7.

24. Bassin RH, Tuttle N, Fischinger PJ. Rapid cell culture assay technic for murine leukaemia viruses. Nature. 1971;229(5286):564–6.

25. Sastry L, Xu Y, Duffy L, Koop S, Jasti A, Roehl H, et al. Product-enhanced reverse transcriptase assay for replication-competent retrovirus and lentivirus detection. Hum Gene Ther. 2005;16(10):1227–36.

26. Stuelke EL, James KS, Kirchherr JL, Allard B, Baker C, Kuruc JD, et al. Measuring the Inducible, Replication-Competent HIV Reservoir Using an Ultra-Sensitive p24 Readout, the Digital ELISA Viral Outgrowth Assay. Front Immunol. 2020;11:1971.

27. Cornetta K, Lin T-Y, Pellin D, Kohn DB. Meeting FDA Guidance recommendations for replication-competent virus and insertional oncogenesis testing. Molecular Therapy Methods & Clinical Development. 2023;28:28–39.

